# Expression of DNA repair genes is modulated during differentiation of olfactory sensory neurons

**DOI:** 10.1101/2023.04.06.535865

**Authors:** Fernanda T. Rowies, Caio M.P.F. Batalha, Thiago S. Nakahara, Bettina Malnic, Nadja C. de Souza-Pinto

## Abstract

Olfactory dysfunction is considered a biomarker of several pathological conditions, including age-associated neurodegenerations, glioblastoma and COVID-19. Olfactory sensory neurons (OSNs) are specialized neurons that detect odorants and send olfactory information to the brain through the olfactory bulb. To perform their function, they are in direct contact with the environment, where they are exposed to several environmental toxins such as atmospheric levels of O_2_ and volatile molecules. Nonetheless, very little is known about DNA damage levels and expression of DNA repair pathways in these cells. Here we measured nuclear and mitochondrial DNA damage in olfactory epithelium (OE) and compared with levels detected in olfactory bulb (OB) and temporal cortex (TC), as a non-olfactory related central nervous system region. Surprisingly, DNA damage was lower in OE and OB when compared with TC, both for nuclear and mitochondrial genomes. Accordingly, expression of representative genes for all excision repair pathways was detected in OSNs. Moreover, expression of most evaluated DNA repair genes was lower in mature versus OSN progenitors, suggesting that DNA repair is downregulated during differentiation. Analysis of single cell expression data confirmed that expression of the most differentially expressed DNA repair genes decreased from progenitor to mature OSNs. Finally, *in situ* hybridization data showed that APE1 mRNA levels are lower in the mature OSNs layer of the olfactory epithelium, closest to the nasal cavity lumen. Altogether, we show here that DNA repair pathways are relevant in protecting OSNs against DNA damage accumulation and that differentiation through the OE is accompanied by changes in the expression levels of DNA repair genes.

## 1. Introduction

Mammals rely on smell for a large variety of physiological and behavioral responses essential to survival, such as detecting prey and predator, possible mates, and food sources. The volatile compounds that make up the various smells are detected by specialized neurons localized in the olfactory epithelium (OE) that covers the nasal cavity, the olfactory sensory neurons (OSNs). OSNs are bipolar neurons with a single dendrite that reaches the OE surface, projecting about 20 to 30 very thin cilia in the nasal mucosa, and whose axon is projected directly to the olfactory bulb (OB) in the central nervous system (CNS). To fulfill their function, OSNs are, necessarily, in permanent contact with the outside environment through olfactory receptors, making them the only neuronal population directly and constantly exposed to air and, thus, to DNA damage agents such as atmospheric O_2_ (Perry et al., 2003) and air pollutants (Valavalandis et al. 2013). Nonetheless, very little is known about DNA damage accumulation and handling in OSNs.

For most regions in the CNS, neurogenesis is restricted to the embryonic and perinatal periods (Cone and Cone, 1976). In adults, this process is restricted to two very specific regions, the lateral subventricular zone (SVZ) and the dentate gyrus of the hippocampus (Kemperman et al., 2018). Neurons in replicative phase are found in the SVZ and migrate, throughout differentiation, to the OB region, which is composed exclusively of differentiated cells (Doetsch *et al*., 1999; Lois and Alvarez-Buylla, 1994). The OE, on the other hand, presents neurogenesis throughout much of adulthood (Schwob et al., 2017), having its neuronal population composed by replicative, immature, and fully differentiated cells, constantly replaced throughout life. In this context, it is believed that the regeneration capacity of OE by neurogenesis is essential for the continuity of olfactory function even after massive damage, as this tissue is capable of not only replace neurons that were lost, but also reestablish the synaptic flux to the correct glomeruli in OB after cellular lesion (Cheung *et al*., 2014).

DNA damage accumulation impairs proliferation, differentiation and affect cellular homeostasis. Even in non-replicative cells, like most CNS neurons, DNA damage impairs cellular fitness and could lead to cell death. Not surprisingly, accumulation of DNA damage has been extensively associated with the pathophysiology of several neurodegenerative diseases, particularly the age-associated neurodegenerations like Parkinson’s, Alzheimer’s, and Huntington’s diseases (Scheijen & Wilson III, 2022). In addition, mutations in DNA repair genes also cause several neurodegenerative diseases, like ataxia telangiectasia (Shiloh, 2020) and other ataxias (Yoon & Caldecott, 2018), amyotrophic lateral sclerosis (Konopka & Atkin, 2022), Cockayne syndrome and some complementation groups of xeroderma pigmentosum (Madabhushi et al., 2014), underpinning the causative link between DNA damage accumulation and neurodegeneration.

Despite the obvious role of genomic stability in neurons, and in contrast to most regions of the central nervous system, very little is known about DNA repair in OSNs. Two relatively recent studies have shown a functional relationship between base excision repair factors and olfactory function. Canugovi and colleagues (Canugovi *et al*., 2015) have shown that mice lacking the DNA glycosylase NEIL1 have impaired olfactory function. NEIL1 expression levels were found to be higher in the olfactory bulb when compared to other brain regions, suggesting a major role in this region. Another study showed that heterozygosity of DNA polymerase β causes neuronal cell death in the OB and impaired olfactory function in an Alzheimer’s disease (AD) mouse model (Misiak et al., 2017). It is noteworthy that olfactory dysfunction has been proposed as a biomarker for early diagnosis of age-associated neurodegenerative diseases such as Alzheimer’s and Parkinson’s (PD), since close to 90% of early-stage AD and PD patients present olfactory dysfunction (reviewed in Dan et al., 2021). Both NEIL1 and DNA pol β are components of the base excision repair pathway, which repairs small base modifications like oxidations and methylations, abasic sites and single strand breaks, which are lesions commonly generated by normal aerobic metabolism (Muftuoglu et al., 2014). Interestingly, Bermudez & Allen (1984) have shown that rat nasal epithelial cells exposed to methyl-methane sulfonate, an agent that induces DNA base methylation, display unscheduled DNA synthesis, indicative of DNA repair. Moreover, Tang and colleagues have recently shown that volatile signals from gamma-irradiated *C. elegans* alleviate embryonic lethality in worms subsequentially irradiated, suggesting that odorants activate a DNA damage response (Tang et al., 2020). However, despite these data suggesting a pivotal role for DNA repair in olfactory function, no study has yet systematically investigated the expression of DNA repair pathways in mature OSNs, or during their maturation processes. In this study, using publicly available gene expression data, we show that OSNs express all major excision repair pathways at levels comparable with molecular pathways relevant to olfactory function. Moreover, expression of DNA repair genes is modulated during differentiation and, surprisingly, olfactory related CNS regions show significantly lower DNA damage levels than a non-olfactory related region, the temporal cortex. Altogether, our data show that DNA repair activities are required for OSN homeostasis and likely contribute to olfactory function.

## 2. Materials and methods

### 2.1. Mice

Newborn (P4 to P7) and 3-week-old (P21-P23) C57BL/6J mice were used in this study. The animals were bred and maintained at the Institute of Chemistry Animal Facility with standard chow and temperature, in a 12 h light/dark cycle. Newborn mice were kept with the mothers up to 1h before euthanasia. Three-week-old mice were euthanized in a CO_2_ chamber, while for newborns a combination of hypothermia and CO_2_ was used. All animal experiments were previously approved by the Institute of Chemistry Animal Care and Use Committee, under protocol # 60/2017, and followed the “Guia Brasileiro de Produção, Manutenção ou Utilização de Animais para Atividades de Ensino ou Pesquisa Científica” (Braga et al., 2019).

### 2.2. DNA isolation

Total DNA was isolated from three adult mouse neuronal regions: OE, OB and an arbitrary “non-olfactory” region of the cortex (CT), using the DNeasy Blood and Tissue (Qiagen, Hilden, Germany) kit, following the manufacturer’s instructions. The amount and purity of the DNA samples were measured using a Nanodrop® (Thermo Scientific, Waltham, MA). Samples were diluted to a 6 ng/μL in DNAse-free water before amplification reactions.

### 2.3. Quantification of DNA lesions by long extension PCR assay

Nuclear and mitochondrial DNA damage levels were measured by a long extension PCR assay adapted from Kovalenko and Santos, 2009. DNA lesion frequency was obtained from the amplification ratio of a long and a short amplicon for each genome (Fig. 1A). Primer sequences and experimental conditions for each reaction and amplification products are summarized in Supplementary Table S1. For amplification of the long amplicons, *AccuPrime Tak DNA Polymerase High Fidelity* (Invitrogen) was used, while for the short we used the *Taq DNA Polymerase Recombinant* (Invitrogen) kit, following manufacturer’s instructions. Experimental conditions for both amplification reactions are presented at Supplementary Table S2.

**Figure 1:**
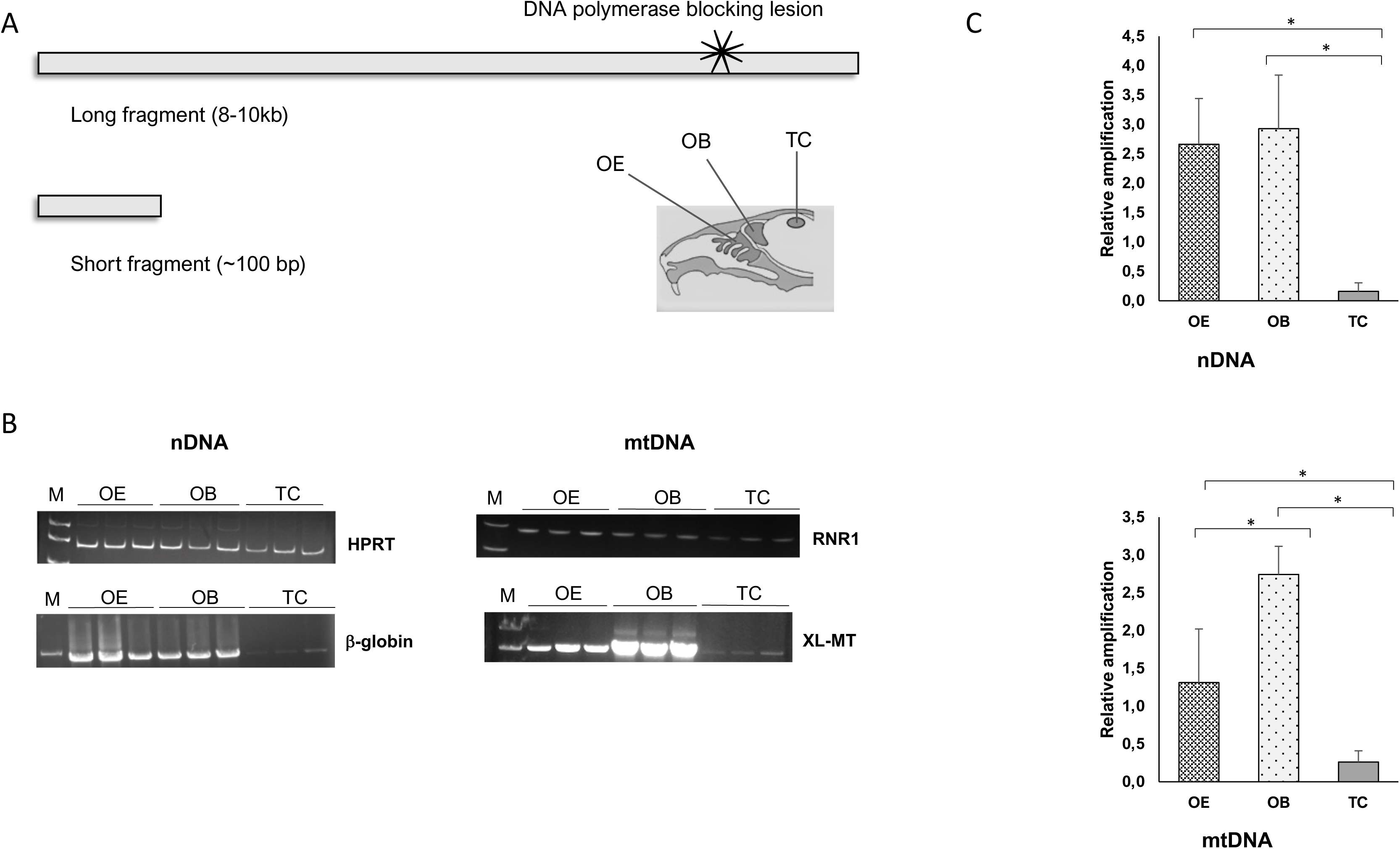
Quantification of nuclear (nDNA) and mitochondrial (mtDNA) DNA damage in olfactory epithelium (OE), olfactory bulb (OB), and temporal cortex (TC) from 3-week-old mice. **A**) Cartoon depicting the different amplicons with a lesion (X) and the anatomical regions used for analyses. **B**) Representative images of the amplicons resolved in 1% agarose (long fragments, lower panels) or 10% PAGE (short fragments, upper panels) gels. **C**) Relative amplification of the long fragments normalized by amplification efficiency of the short fragments in the three tissues. Values are average ± SD of three independent experiments, performed in duplicate. * Indicates p < 0.05, as calculated by *One-way ANOVA*.

Amplification products were resolved by electrophoresis; long fragments in a 1% agarose gel in TAE (40 mM Tris, 20 mM acetic acid, 1 mM EDTA, pH 8.0) buffer, and short fragments in a 10% PAGE (19:1, acrylamide:bis-acrylamide) gel in TBE (89 mM Tris, 89 mM boric acid, 2 mM EDTA, pH 8.0) buffer. Following electrophoresis, gels were incubated in ethidium bromide for 20 min, distained in water for 10 min and photographed under UV light. Band intensity was obtained using ImageJ vl.48 software.

To calculate DNA damage levels, we normalized amplification of the long fragment by the short fragment (A_long/short_) in each group (OE, OB and TC) to obtain corrected amplification values (A_OE_, A_OB_ and A_TC_), as described in Kovalenko & Santos, 2009. Data presented are the corrected relative amplification for each region.

### 2.4. Differential expression analysis with simulated dispersion values

Fastq data reported in Magklara et al., 2011 was obtained from the authors upon request, and aligned to the m39 mouse genome build using bowtie2 and Tophat2 (Langmead B and Salzberg SL, 2012; Kim et al., 2013). A read count matrix was then constructed using the featureCounts function of Subread for both ngn and omp samples (Liao et al., 2013; Liao et al., 2014). The resulting raw read count matrices were further analyzed in R (Team RC, 2022). Ensembl gene id was converted to gene symbols using the biomaRt package, and the data was analyzed using the edgeR package (Chen et al., 2016; McCarthy et al., 2012; Robinson et al., 2010). Since the data did not have biological replicates, it was not possible to carry out an adequate differential expression analysis. However, due to the scarcity of data suitable for our analysis in the literature, we decided to analyze the data using a simulated dispersion value, following recommendations regarding the use of data without replicates in the edgeR user guide. We chose a conservative BCV (square-root-dispersion) of 0.4 and simulated differential expression using the exactTest function. Genes were considered differentially expressed with an adjusted p-value (FDR) < 0.05. Finally, we used biomaRt (Durinck et al., 2005; Durinck et al., 2009) to filter genes associated with the GO id GO:0006281 (DNA repair). Simulating differential expression with a dispersion value is in no way a substitute for biological replication, and the results should be interpreted with extreme caution, and in conjunction with other more robust results.

### 2.5. Single-cell expression trajectory analysis

Single-cell gene expression data of nasal olfactory neurons in several stages of mice olfactory neurogenesis were obtained from Hanchate et al., 2015. Single-cell trajectory analysis was then carried out using the Monocle R package, according to the user guide (Trapnell et al., 2014; Qiu et al., 2017; Qiu et al., 2017). Genes were chosen as markers for each stage of the neuronal differentiation were: Ascl1 (progenitor); Neurog1 and/or Neurod1 (precursor); Gap43 and/or Gng8 (immature OSN); Omp and four olfactory sensory transduction molecules downstream of odorant receptors – Gnal, Adcy3, Cnga2 and Cnga4 (mature OSN). Cells were then ordered and plotted in pseudotime, and differential gene expression was calculated using the differentialGeneTest function. Differentially expressed genes along the pseudotime trajectory were considered using an adjusted p-value (FDR) < 0.05. DNA repair genes (GO:0006281) were filtered. We then selected the same 12 genes that were found to be differentially expressed in the differential expression analysis of the Magklara et al., 2011 dataset with simulated dispersion values and plotted their pseudotime expression for comparison with the previous results, including those that had an adjusted p-value (FDR) > 0.05. Additionally, we clustered the pseudotemporal expression pattern of all genes differentially expressed in this analysis into 3 clusters, using the plot_pseudotime_heatmap function.

### 2.6. Gene ontology enrichment analysis

We enriched the gene sets that composed the 3 clusters found in the pseudotime trajectory analysis for GO biological process terms using the shinyGO tool (bioinformatics.sdstate.edu/go/), version 0.76 (Ge et al., 2020). We considered terms enriched with adjusted p-value (FDR) < 0.05.

### 2.7. OE dissection and sample preparation

3-week-old mice were euthanized as described, and whole OE and OB tissues were dissected under a stereoscopic microscope. All utensils were cleaned with RNAse inhibitor solution. For RNA extraction, tissue samples were flash-frozen in liquid N_2_, while for *in situ* hybridization the organs were immediately put in paraformaldehyde 4% and left overnight at 4°C. Then, they were transferred to RNAse free PBS containing 0.45 M EDTA and left overnight. After that, the samples were transferred to PBS containing 20% (m/v) sucrose and incubated for 30 minutes. The samples were then carefully removed and wiped, transferred to plastic containers with Tissue Tek® O.C.T. Compound (Sakura Finetek, Torrance, CA) and stored at -80°C.

### 2.8. Total RNA extraction and cDNA synthesis

RNA from whole mouse OE samples was extracted using RNeasy kit (Qiagen, Hilden, Germany), following the manufacturer’s instructions. The amount and purity of RNA were checked spectrophotometrically in a Nanodrop® (Thermo Scientific, Waltham, MA), while integrity (18S and 28S bands) was checked by electrophoresis in a 1% agarose gel in TAE (40 mM Tris; 20 mM acetic acid, 1 mM EDTA, pH 8.0) containing 1% (v/v) commercial household bleach (Aranda et al, 2013). Complementary DNA (cDNA) was obtained from total RNA using *High-Capacity cDNA Reverse Transcription kit* (Thermo Scientific, Waltham, MA) following the manufacturer’s instructions, in 25 μL reactions using up to 2 μg of total RNA and Oligo d(T) primers.

### 2.9. Gene expression analysis

Expression level of genes of interest in OE cDNA was measured by quantitative real-time PCR (qRT-PCR), in 20 μL triplicate reactions using *Power SYBR Green PCR Master Mix* (Thermo Scientific, Waltham, MA), according to manufacturer’s instructions. Expression of Hydroxymethylbilane synthase (HMBS, gene ID 15288) was used as housekeeping control. Primer pairs for the DNA repair genes were designed using *PrimerBank* (Wang & Seed, 2003) and validated by *BLAST*. Primer sequences are presented in Supplementary Table S3. All primer pairs were validated (Supplementary Table S4) before experiments. Amplification reactions, containing 0.72, 0.36, 18 or 90 ng of cDNA, were carried out in a 7300 Real Time PCR System (Thermo Scientific, Waltham, MA). Relative expression was calculated according to Pfaffl (2001) with adaptations, using the amplification efficiency (Supplementary Table S4) of the housekeeping gene and genes of interest (AF_hkg_ and AF_int_, respectively) and Δ^ct^ values for newborn and 3 weeks old mice (⊗^ct nwb^ and ⊗^ct 3wo^, respectively), as follows:

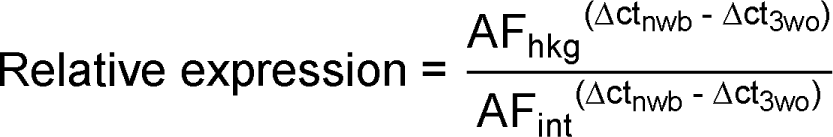

### 2.10. RNA probe

APE1 cRNA probe was *in vitro* translated using digoxigenin-labeled uracil as described in Camargo et al 2019. For that, the genomic sequence was obtained from GeneBank and used to design primers to amplify ∼1000 pb of the coding sequence of the mRNA using GENtle (http://gentle.magnusmanske.de/). Inserts (primers and size are presented in Supplementary Table S5) were cloned into pGEM-T Easy (Promega, Madison, WI) vector and confirmed by restriction and sequencing analysis. Plasmids containing correct sequences were linearized so that the inserts were contiguous to the SP6 or T7 promoters and used for the *in vitro* translation reactions.

### 2.11. *In situ* hybridization

*In situ* hybridization was carried out as described in Nakahara et al., 2014 and Ishii et al., 2004. Briefly, slides were fixed with paraformaldehyde 4% for 20 min, washed and treated with 0.1 M HCl for 10 min and with 0.1% (v/v) H_2_O_2_ for 30 min. Samples were acetylated with 1 mL of acetic anhydride in 250 mL of 0.1 M triethanolamine pH 8.0, and hybridized with pre-heated hybridization solution (50% (v/v) formamide, 10% (m/v) dextran sulfate, NaCl 0.6 M, yeast tRNA 200μg/ml, 0.25% (m/v) SDS, Tris-HCl 10 μmM pH 8.0, 1X Denhardt solution, EDTA 1mM), containing sense and antisense RNA probes, for 16 h at 60°C, covered by glass coverslip. On the next day, coverslips were displaced in 5X SSC (1X SSC: 150 mM NaCl, 15 mM sodium citrate, pH 7.0) and washed at 60°C in 2X SSC for 30 min, in 0.2X SSC for 20 min and 0.1X SSC for 20 min. Coverslips were transferred to 0.1X SSC solution at room temperature, and then subsequentially to: i) PBS containing 0.1% (v/v) Tween-20 (ThermoFisher Scientific, Waltham, MA) for 10 min; ii) TN (100 mM Tris-HCl pH 7.5, 150 mM NaCl) buffer for 5 min, twice. Coverslips were blocked with TNB (100 mM Tris-HCl pH 7.5, 150 mM NaCl, 0.05% PerkinElmer FP1020 [Waltham, MA]) buffer, then incubated with antibody anti-DIG-AP (Roche, Basel, Switzerland), diluted 1:800 in μ200 L TNB, covered with Parafilm™ (Sigma-Aldrich, St. Louis, MO), at 4°C overnight.

After incubation, coverslips were washed 6 times in TNT (100 mM Tris-HCl pH 7.5, 150 mM NaCl, 0.05% Tween-20) buffer for 5 min each, with gentle agitation, washed twice in Alkaline Phosphatase buffer (100 mM Tris-HCl pH 9.5, 50 mM MgCl_2_, 100 mM NaCl, 0.1% Tween-20) for 5 minutes and incubated with a solution of 1:1 (v/v) 2X Alkaline Phosphatase buffer (10% (v/v) Polyvinyl Alcohol (Mowiol, Sigma-Aldrich, St. Louis, MO), previously filtrated and added 6.7 mL of 75 mg/mL NBT and 5 mL de 50 mg/mL BCIP). Mounted coverslips were visualized using a Nikon microscope, under 4 and 10x magnification.

### 2.10. Statistical analyses

Experimental results are presented as mean ± standard deviation, except when otherwise indicated. Statistical analyses were carried out using Excel software (Microsoft) by *One-Way* ANOVA, assuming p < 0.05 as statistically different.

## 3. Results

### 3.1. Olfactory-associated tissue accumulate less DNA damage than CNS tissue

Olfactory sensory neurons are the only neuronal cells that are in direct contact with the environment. Consequently, mature OSNs are exposed to exogenous genotoxins such as environmental pollutants and atmospheric O_2_ tension, in addition to the endogenously generated genotoxins other neuronal cell types are exposed. Thus, we hypothesized that OSNs accumulate more DNA damage than other neuronal cells. To address this question, we used a long extension PCR assay to measure nuclear and mitochondrial DNA damage in total DNA obtained from murine OE in comparison to OB and to temporal cortex (TC), which was included as a control CNS region not involved in the olfactory system. The assay measures amplification efficiency of a long DNA fragment (∼ 10kb), taking advantage of the fact that the presence of DNA lesions will block the progression of the DNA polymerase. Thus, lower amplification efficiency indicates presence of more lesions in the template. Amplification of the long fragment is normalized by the amplification of a short amplicon of the same region (Fig. 1A), considering that the chances of harboring a blocking DNA lesion is much higher in the long fragment, the amplification of the short fragment functions as normalization for DNA content in the reaction.

Using this assay, we quantified amplification of a 10Kb fragment of the β-globin gene, to assess DNA lesion in the nuclear DNA, and of a 16Kb fragment of the mitochondrial DNA, comprising almost the entire mouse mitochondrial genome. These amplifications were normalized by short amplicons of the HPRT gene and the RNR1 gene, respectively. For each brain region and amplification target we used 3 biological samples. Figure 1B shows a representative image of the amplification products, and figure 1C shows the quantification of the results, expressed as normalized amplification for each independent sample. Surprisingly, in both nuclear and mitochondrial DNA, TC showed significantly lower amplification efficiency than the other two regions, the olfactory related tissues OE and OB, suggesting that the steady-state levels of DNA lesions are lower in olfactory related neuron, when compared to a non-related region of the CNS. In mtDNA, but not in nDNA, OE had significantly lower amplification than OB, suggesting more DNA lesions in mtDNA in these cells.

The observation that OE and OB have lower steady-state levels of DNA lesions than TC prompted the question whether these cells are DNA repair proficient, as there is no systematic characterization of DNA repair in olfactory neurons, and to our knowledge, only three articles have investigated DNA repair in the context of olfactory tissues (Bermudez & Allen, 1944, Canugovi *et al*., 2015; Misiak *et al*., 2018).

### 3.2. OSNs express all major DNA repair pathways

To begin to address that, we manually extracted expression values, in fragments per kilobase of exon per million mapped fragments (FPKM), of representative DNA repair genes in two published mouse OE transcriptomes, one comparing progenitor and mature OSNs (Magklara et al., 2011) and another comparing OE from neonates (enriched in progenitor cells) and 4-week-old mice (enriched in mature OSNs) (Camargo et al., 2019). We curated data from 4 representative genes of 4 of the 5 excision repair pathways, base excision (BER) and nucleotide excision (NER) repair, homologous recombination (HR) and non-homologous end joining (NHEJ). The mismatch repair pathway was not included here as mature OSNs are terminally differentiated. The FPKM values obtained for each gene in the two different transcriptomes are presented in Table 1. With few exceptions, like EXO1 and UNG in mature (OMP^+^) OSNs, all genes have detectable expression.

**Table 1:**
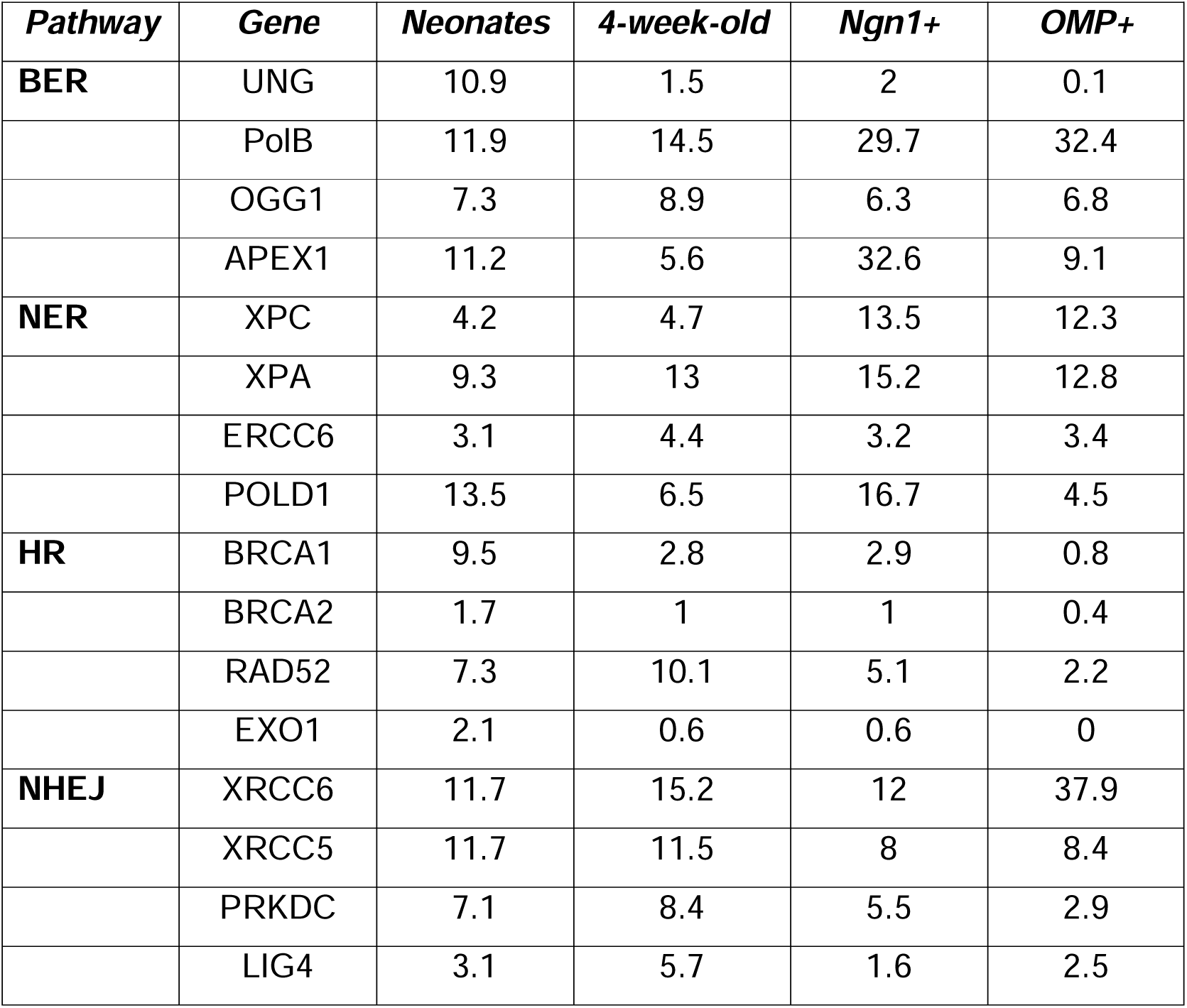
FPKM (fragments per kilobase of exon per million mapped fragments) values for selected DNA repair genes, manually extracted from the data published in Camargo et al., 2019 (neonates/4-week-old) or Magklara et al., 2011 (Ngn1+/OMP+).

And although the absolute values are somewhat different between the two datasets, most genes showed similar pattern of change in expression when comparing neonates/Ngn1^+^ cells to 4-week-old/OMP^+^ cells. In addition, the expression levels for most DNA repair genes are comparable to that of neuronally-relevant pathways, like Rho GTPase axonal orientation and Ca^2+^ signaling in synapses. In the dataset published by Camargo et al., 2019, the expression range for 4 genes of these two pathways (CREB1 and GRIN2A for Ca^2+^ signaling, and CXCR4 and DBN1 for axonal orientation) were within the range 1.5 – 27.1 for neonates and 1.2 – 11.3 for 4-week-old mice. For comparison, for all DNA repair genes analyzed here, the expression range was 1.7 – 13.5 for neonates and 0.6 – 15.2 for 4-week-old, suggesting that DNA repair genes are expressed at relevant levels in OSNs of the OE. Moreover, for most genes analyzed, there was a trend to lower expression in mature versus progenitor populations.

We validated these findings using RT-PCR with RNA extracted from total OE obtained from newborn or 3-week-old mice, again with the rationale that tissue from newborn mice are enriched in progenitor cells compared with the 3-week-old, which are enriched in mature OSNs. This assumption was validated by measuring expression of the OSN differentiation markers OMP, Ngn1 and Ric-8b (Fig. 2A). Relative expression of the 16 DNA repair genes is shown in Fig. 2B. For almost all genes evaluated, with exception of the NER proteins XPC and POLD, expression levels were significantly decreased in 3-week-old OE compared with neonates, suggesting that DNA repair pathways are actively expressed in OSNs and that expression of DNA repair genes is modulated by differentiation.

**Figure 2:**
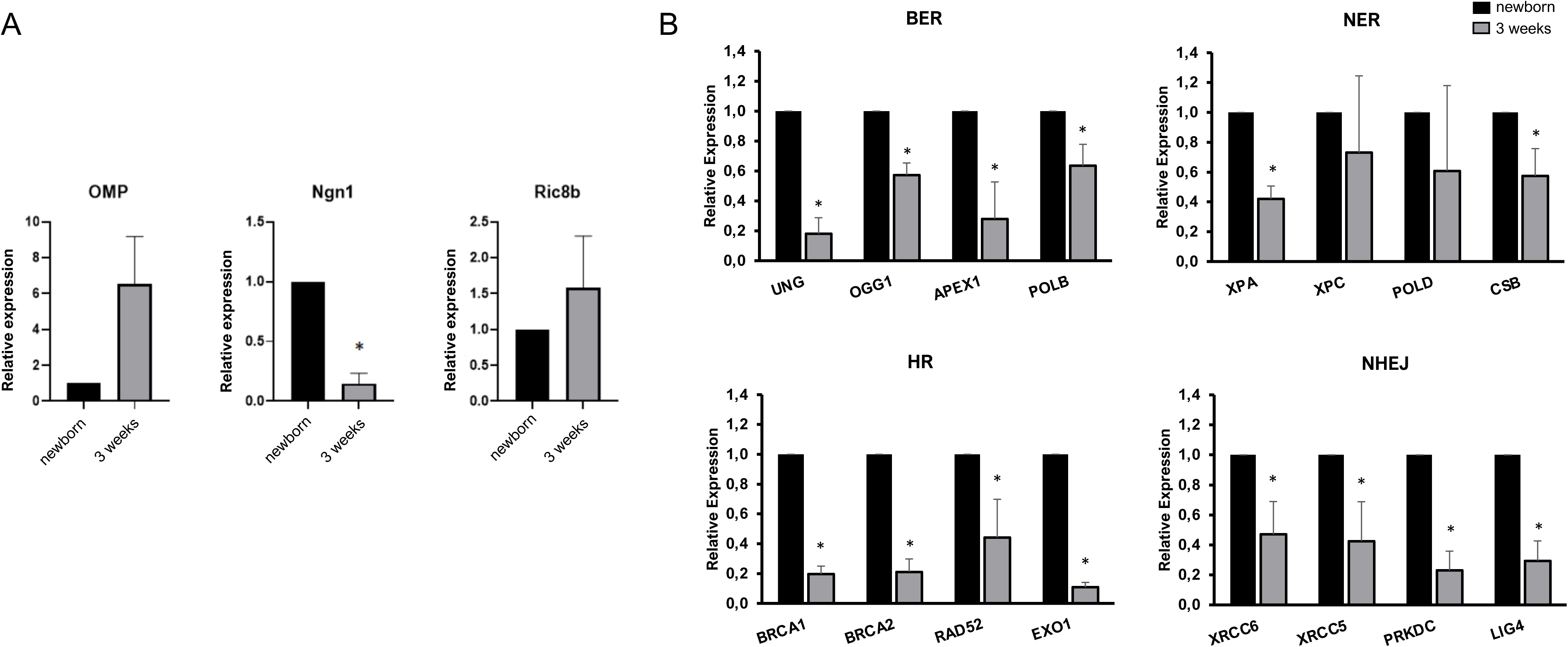
Analysis of differential expression of DNA repair genes in mouse OE. OE was dissected from neonatal and 3-week-old mice and gene expression was analyzed by RT-PCR, as described. **A**) Expression of markers of OSN differentiation, Ngn1 (progenitors), OMP and Ric-8b (mature OSNs); **B**) Expression of the representative genes of the BER, NER, HR and NHEJ pathways found to be differentially expressed in the manually curated analysis shown in Table 1. Relative expression was calculated as proposed by Pfaffle, 2001. Values are average ± SD of two independent experiments, performed in triplicate. * Indicates p < 0.05, as calculated by *One-way ANOVA*.

### 3.3 Expression of DNA repair genes is modulated during OSN differentiation

To test the hypothesis that DNA repair is modulated during OSN differentiation we first analyzed all differentially expressed genes, as logFC, between Ngn1 and OMP-positive populations from the Magklara et al., 2011 study, using the edgeR R package. Unfortunately, this dataset contains no replicates for the two populations, which makes it impossible to carry out an adequate differential expression analysis. Nonetheless, due to the scarcity of proper datasets compatible with our study, we decided to analyze this dataset using the simulated dispersion values (see the methods section). While nothing can substitute biological replication, we believe that when the results of the differential expression with simulated dispersion are analyzed in a conservative manner and in conjunction with the other results in this study, they corroborate the overall results of our paper. DNA repair genes were filtered from the dataset of differentially expressed genes. Figure 3A lists 12 differentially expressed DNA repair genes, with logFC higher/lower than 2. From the 12 genes listed, 11 were downregulated during differentiation and only one, Polk, was upregulated from Ngn1^+^ to OMP^+^ cells.

**Figure 3:**
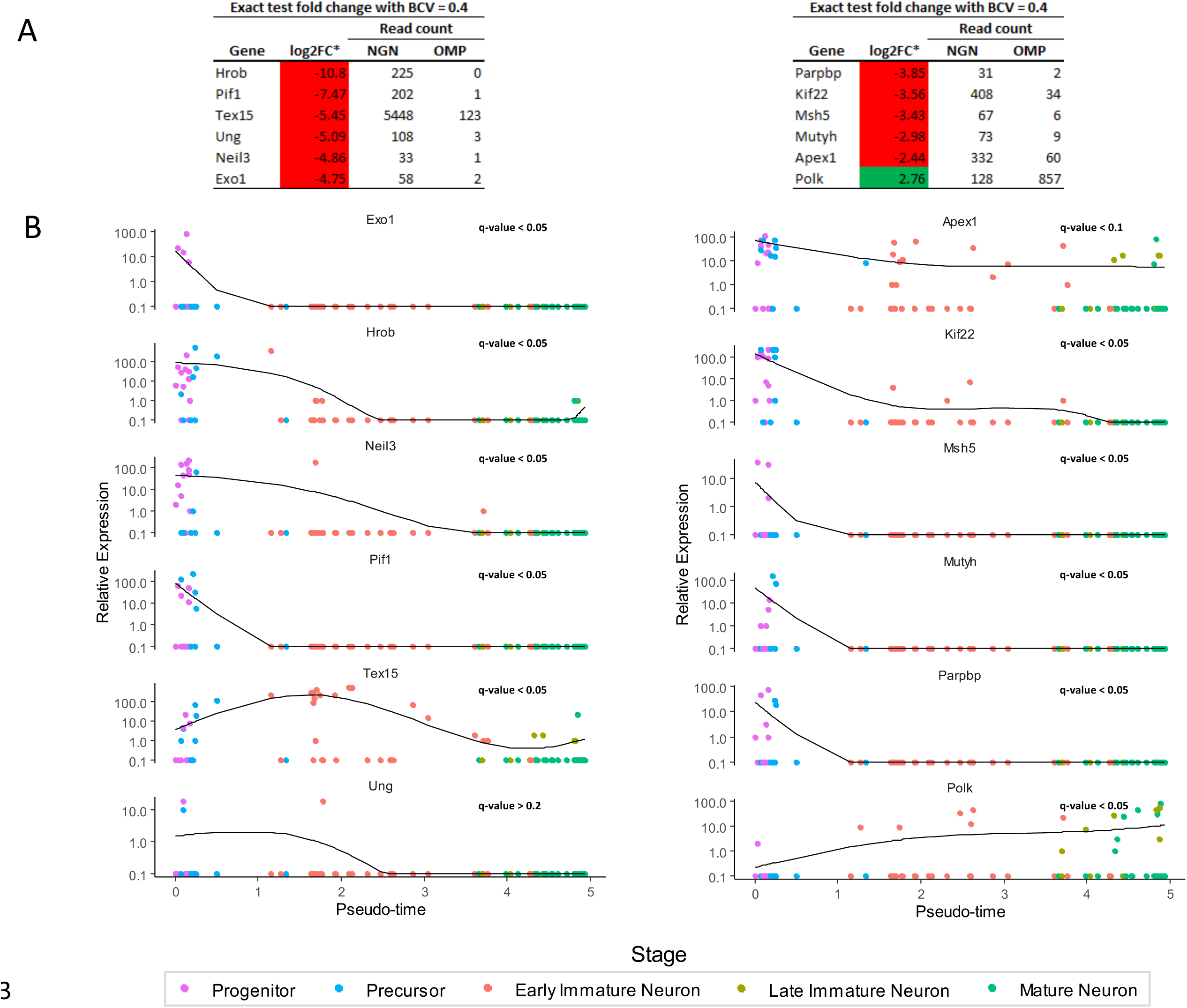
Analysis of differential expression of DNA repair genes in OE. **A**) Difference in expression of DNA repair genes, expressed as Log 2-fold change (Log2FC) using the dataset from Magklara et al. (2011). Since the two samples have no replicates, a true differential expression statistic cannot be calculated. Instead, we used a simulated biological coefficient of variation to identify potential differentially expressed genes. The tables show the 12 genes that were significantly differentially expressed in this simulation in mature OSNs (OMP^+^) versus intermediate progenitors (Ngn1^+^) with a biological coefficient of variation (BCV) of 0.4. **B**) Pseudo-time trajectory analysis of expression of the DNA repair genes potentially differentially expressed, shown in A, in the single-cell RNA-Seq data from Hanchate et al. (2015), with accompanying q-value (FDR corrected p-values). *the exactTest function of edgeR shinks the Log2FC, thus preventing infinite log-fold-changes, as would be the case for genes with zero counts in one of the samples.

To further investigate this relationship, we analyzed the data from Hanchate et al., 2015, who performed single cell RNA transcriptomics in progenitor, precursor, immature and mature OSNs. Several genes were found to be differentially expressed in an expression pseudotime trajectory analysis: 96 using a q-value of 0.05 and 123 using a q-value of 0.1, out of a total of 481 DNA repair genes. The full results for the expression trajectory analysis can be found in Supplemental Table S6. For all the 12 differentially expressed DNA repair genes listed in Fig 3A, the expression trajectory along pseudo-time is shown in Figure 3B. There is striking similarity between the logFC changes seen between Ngn1^+^ and OMP^+^ populations in the Magklara et al. dataset and the pseudo-time trajectory of the same genes in the single cell analysis from Hanchate et al., suggesting that the expression of, at least this set of DNA repair genes, is downregulated during OSN differentiation. Only Apex1 and Ung did not display a q-value of less than 0.05 in the expression trajectory analysis. Apex1, however, did display a q-value lower than 0.1, being considered significant using a more relaxed threshold, while Ung displayed a q-value of ∼0.205. Both, however, were found to be downregulated in Fig. 2B.

To try to find expression patterns in DNA repair genes during OSN differentiation, all differentially expressed DNA repair genes (considering a q-value < 0.05) in the Hanchate et al. study were clustered according to their expression trajectories along the pseudo-time. The heatmap presented in Fig. 4A shows that DNA repair genes can be organized in 3 clusters. Cluster 3 (blue, left side) groups genes that were upregulated during differentiation, while clusters 1 (green, in the middle) and 2 (salmon, right) group genes that are downregulated, with cluster 1 being the set with the stronger changes. Most genes are clustered within clusters 1 and 2, again suggesting that DNA repair is downregulated during OSN maturation. In total, out of the 96 differentially expressed DNA repair genes at q-value < 0.05, 88 could be grouped in these 3 clusters, with the remaining 8 not displaying an expression pattern compatible with any clustering.

**Figure 4:**
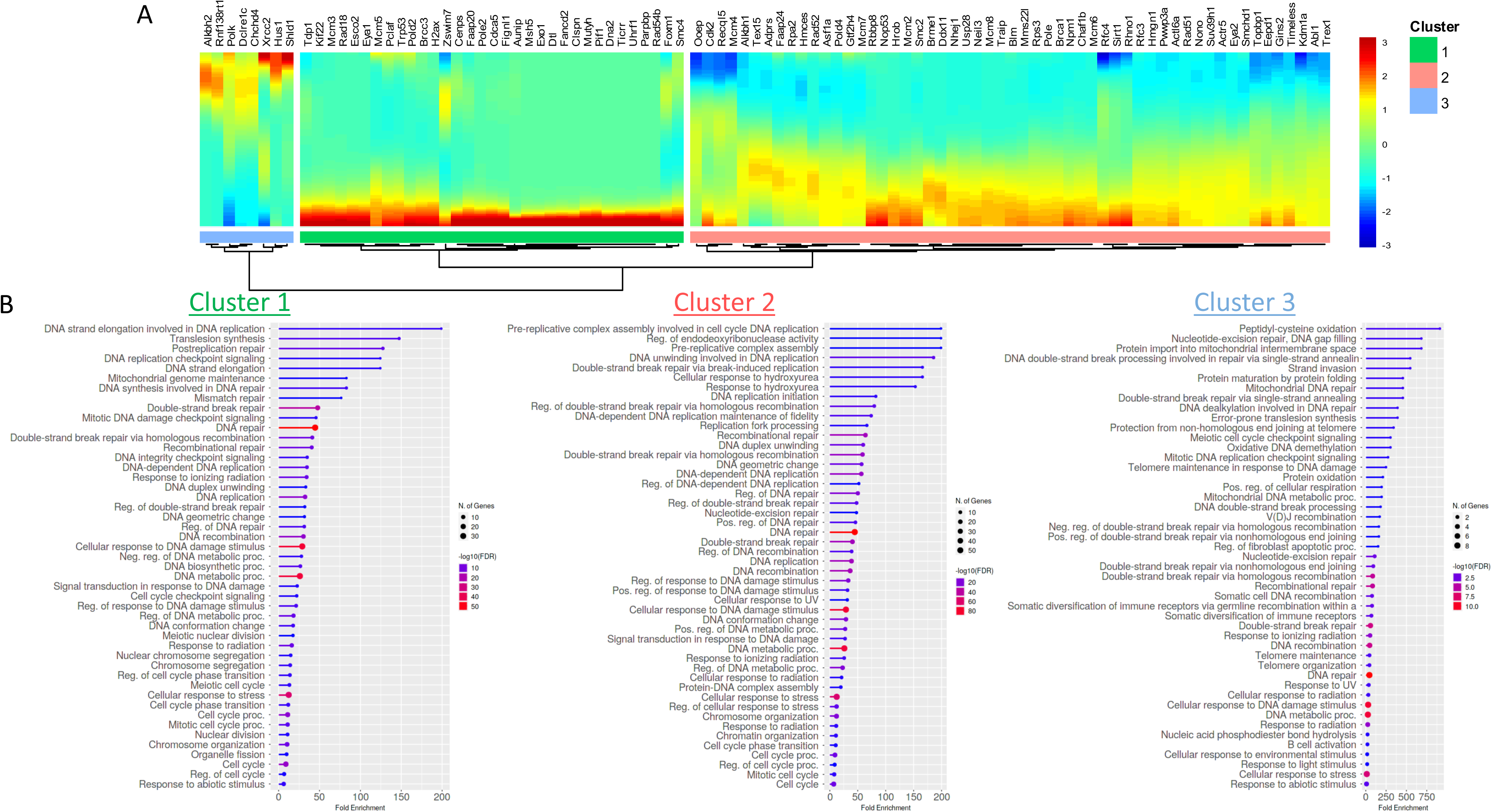
Clustering and GO enrichment analysis of DNA repair genes. **A**) Clustering of DNA repair genes according to their trajectory pattern in the pseudo-time trajectory analysis with single-cell RNA-Seq data from Hanchate et al. (2015). **B**) Top 50 GO biological process terms enriched in the three clusters in A, with FDR < 0.05.

Not surprisingly, the top 10 GO (geneontology.org) biological processes enriched in clusters 1, 2 and 3 are related to DNA repair and DNA replication (Fig. 4B).

### 3.4. APE1 expression is downregulated during OSN migration through the OE

since the transcriptomics analyses suggested that DNA repair genes are downregulated during OSN maturation we tested the hypothesis *in vivo* measuring Apex1 mRNA levels by *in situ* hybridization. The Apex1 gene product, APE1, is the major abasic site endonuclease in BER and is an essential gene in mice, as its knockout is embryonically lethal (Xanthoudakis et al., 1996; Ludwig et al., 1998; Meira et al., 2001). In addition to the DNA repair function, APE1 has an additional function in gene expression regulation, and both functions are essential for cell survival (Izumi et al., 2005). Moreover, Apex1 downregulation in differentiated cells was found in both the Magklara et al. and the Hanchate et al. data sets (Figs. 3A & B) and was validated by RT-PCR in mouse OE (Fig. 2B). Thus, antisense probes for Apex1 mRNA were generated and used to label OE from 3-week-old mice (Fig. 5) The OE can be divided into 3 major layers according to the dominant cellular populations, the mature OSN layer (region A in Fig. 5B), immature OSN layer (region B) and the most distal, the layer composed by progenitor cells (region C). The staining for Apex1 can be easily detected in the 3 layers and clearly differentiate the OE from the connective tissue bellow. Nonetheless, the staining is progressively decreased from the progenitor cells layer to the mature OSN layer, corroborating that expression of this gene is downregulated as OSN mature and migrate through the OE.

**Figure 5:**
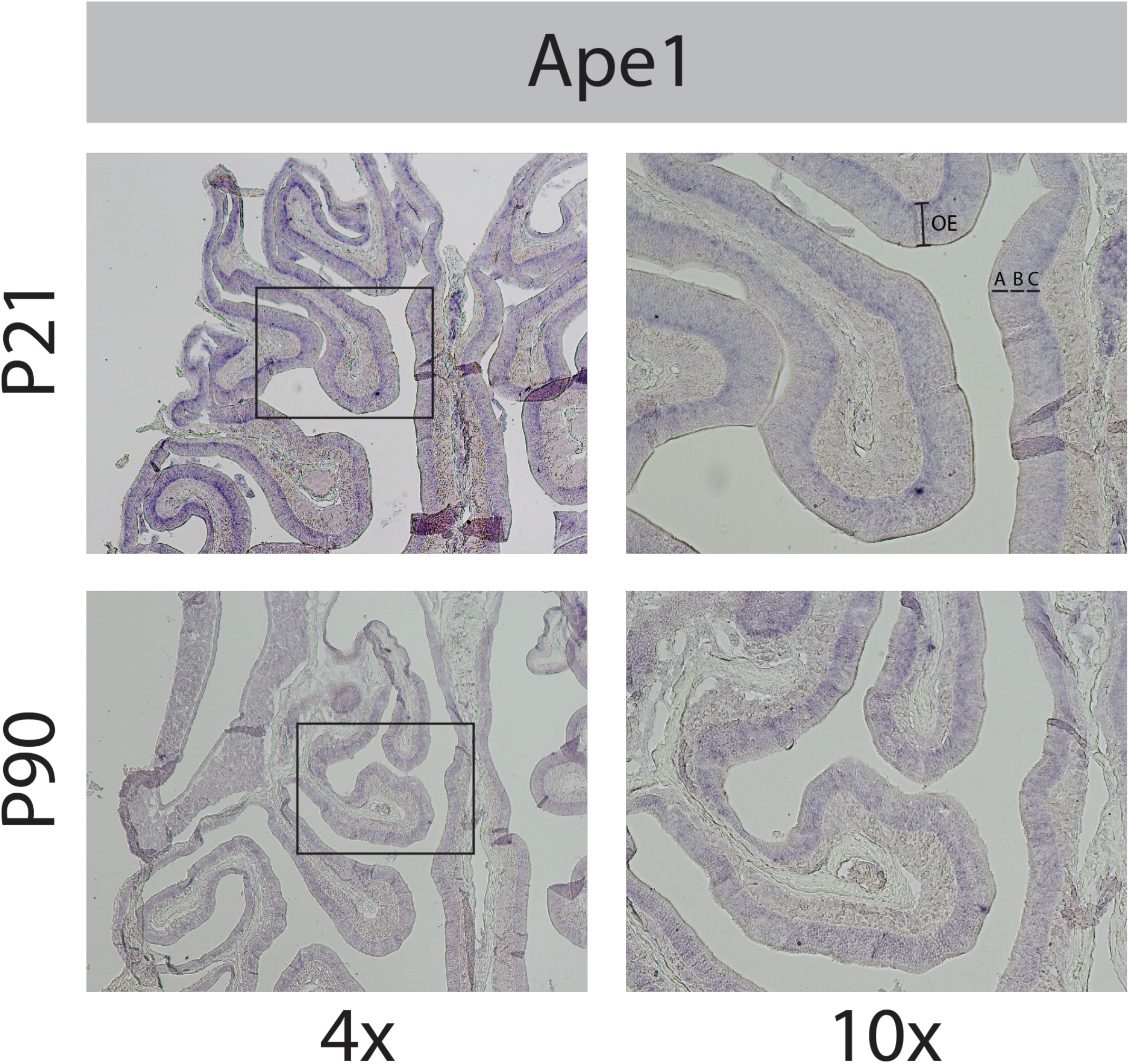
APE1 mRNA levels detected by *in situ* hybridization. Antisense probes for APE1 were used to label OE from 3- and 12-week-old mice. For each age, images at 4 and 10X magnification are shown. (A) mature OSNs layer, (B) immature OSNs layer and (C) progenitor cells. Representative image of three biological replicates.

## 4. Discussion

Genome maintenance is essential for cellular function, even in fully differentiated cells. Neurons are particularly sensitive to genomic instability and several neurodegenerative diseases are caused by defects in DNA repair and aberrant DNA damage response. In the case of OSNs, DNA repair pathways may be even more relevant as these cells are directly exposed to environmental genotoxins and ambient oxygen tension. DNA damage induced by environmental and endogenous conditions has been linked to olfactory dysfunction. For instance, the DNA damaging effects of tobacco components in olfactory mucosa have been long identified (Belinsky et al., 1987). In fact, DNA adducts were detected in nasal olfactory mucosa of rats treated with tobacco components, even when exposed to the drugs in drinking water (Zhang et al., 2009). Increased DNA damage, as measured by the Comet assay, was also detected in olfactory epithelia of dogs living in the city of São Paulo, Brazil, when compared to controls, suggesting that air pollution is an important genotoxin to olfactory cells (Kimura et al., 2010). Moreover, the Harlequin mutation, which downregulates the apoptosis inducing factor (Klein et al., 2002), has been shown to cause olfactory dysfunction in mice through increased oxidative stress and oxidative DNA damage (Vaishnav et al., 2008). Thus, it is reasonable to speculate that DNA repair activities play a relevant role in protecting OSNs homeostasis and olfactory function. In fact, a recent report showed that Sirt1 activity prevents accumulation of DNA double strand breaks in subventricular zone derived stem cells, and that this contributes to maintaining olfactory function in aging mice (Ren et al., 2022). Nonetheless, very little is known about DNA damage and repair in these specialized neurons.

We hypothesized that OSNs accumulated more DNA damage than other neuronal populations due to their fully differentiated status and constant exposure to airborne genotoxins. Surprisingly, using a PCR-based assay that detects polymerase-blocking DNA lesions we found that OE and OB have significantly lower DNA damage, both in nuclear and mitochondrial genomes, than a non-olfactory related region of the brain (Fig. 1). Steady state DNA damage levels result from the balance between formation rate and repair. While basal levels of 8-oxodG in whole brain DNA were comparable to other tissues in rats and mice (Hamilton et al., 2001), levels of DNA single-strand breaks in whole genome were higher in mouse brain than heart, liver, sperm, and bone marrow (Cai et al., 2021). On the other hand, BER activities in mouse brain were not significantly different than levels found in liver and heart (Karahalil et al., 2002), suggesting that brain tissue, in fact, accumulates more DNA damage. Moreover, recent results have suggested that DNA damage, specifically DNA double-strand breaks, are formed because of normal brain activity and are involved in neural plasticity (reviewed in Konopka & Atkin, 2022), and that dysfunction and neurodegeneration arise when there is an imbalance between damage formation and repair. These results underly the role of DNA repair in neuronal cells and reinforce the need for a better understanding of DNA repair pathways in OSNs.

In line with that, we found that representative genes of four major excision repair pathways, BER, NER, HR and NHEJ, are expressed in murine OE, with transcript levels compared to that of genes of other relevant pathways, such as calcium signaling and axonal orientation (Table 1), showing that DNA repair is constitutively expressed in OSNs and must have a relevant functional role. Mismatch repair activities were not evaluated because this repair pathway is physically and functionally associated with the replication fork (Modrich, 2016). It should be noted that similar results were obtained from two independent sets of transcriptome data (Magklara et al., 2011; Camargo et al., 2019), strengthening the conclusion. Moreover, qRT-PCR confirmed the expression of these transcripts (Figure 2).

In most mammalian cell types, DNA repair activity is affected by the differentiation state. This effect was first demonstrated in proadipocytes (Tofilon & Meyn, 1988), and it was later shown that, in general, pluripotent cells have higher DNA repair capacity than differentiated lineages (reviewed in Mani et al., 2020; Sjakste & Riekstiņa, 2021). We have demonstrated earlier that embryonic stem cells have higher DNA repair capacity than various differentiated cell types (Maynard et al., 2008). These findings were not surprising, as terminally differentiated cells do not replicate, which would ease the pressure on genomic integrity maintenance. Nonetheless, differentiated cells are still transcriptionally active and cannot dismiss DNA repair completely. Conversely, several observations also indicate that DNA damage response regulates cellular differentiation (Sherman et al., 2011).

Here we found that OSNs maturation is associated with decrease in expression of DNA repair genes, when comparing whole tissue expression by RT-PCR form neonate to 3-week-old mice (Fig. 2) or in single-cell RNA-Seq data from progenitor to mature neurons (Fig. 3). In the pseudo-time trajectory analysis from the Hanchate et al. data, only eight show higher expression in the differentiated lineages. Moreover, most DNA repair genes in this analysis group in clusters that are downregulated with differentiation, with only few (8) clustering with an upregulation trajectory (Fig. 4).

These results are in line with other observations for neuronal cells. Sykora et al. (2013) showed that undifferentiated human SH-SY5Y neuroblastoma cells are less sensitive to oxidative stress than their differentiated counterparts, in part due to higher BER capacity. Interestingly, they find lower APE1 protein levels in the differentiated versus the undifferentiated cells (Sykora et al., 2013), suggesting the BER gene expression is modulated during the *in vitro* differentiation process. Using *in situ* hybridization to assess mRNA levels we find that APE1 expression levels change in the mouse EO from the progenitor (higher) to the mature OSN layer (lower) (Fig. 5), in correlation with the RT-PCR results shown in Fig. 2B and with the gene expression analysis results in Fig. 3A and 3B. Together, these results indicate that as OSNs mature and migrate toward the nasal cavity lumen, DNA repair is downregulated but not abrogated, as genomic integrity is still required for cellular function.

This is the first direct demonstration of the expression of DNA repair genes in the olfactory epithelium, and together with the few reports on olfactory dysfunction in mice defective in DNA repair enzymes, underscore the need for further investigation on the role of DNA repair activities in maintaining OSNs homeostasis and olfactory function. Considering that olfactory function has been proposed as an independent prognosis factor in glioblastoma (Kebir et al., 2020) and neurodegeneration (Dan et al., 2021), and is a prevalent symptom of several viral infection like COVID-19 (Parma et al., 2020), a better understanding of the molecular mechanisms involved in maintaining OSNs homeostasis may have important implications in human health.

**CRediT authorship contribution statement**: NCS-P, BM & FTR conceived and designed the project. FTR, CMPFB & TN performed experiments. FTR & NCS-P wrote the paper. All authors revised and approved the manuscript.

**Conflict of interest**: The authors declare that they have no conflict of interest with the contents of this article.

## Supporting information

supplemental table 6 with list of genes

## Acknowledgements

**Acknowledgements**: This work was supported by FAPESP grants 2010/51906-1 and 2017/04372-0 to NCS-P and 2016/24471-0 to BM. FTR was supported by FAPESP fellowship 2017/13723-0, CMPFB was supported by CNPq fellowship 140207/2018-0 and TSPN was supported by FAPESP fellowship 2018/11860-4. We thank Dr. S. Lomvardas for access to raw data for their transcriptomic analysis published in Magklara et al., 2011.

## References

Altman J, Das GD. Autoradiographic and Histological Evidence of Postnatal Hippocampal Neurogenesis in Rats. The Journal of Comparative Neurology, 124(3): 319–335; 1965.

Aranda PS, LaJoie DM, Jorcyk CL. Bleach gel: a simple agarose gel for analyzing RNA quality. Electrophoresis, 33 (2): 366–9, 2012.

Belinsky SA, White CM. Devereux TR, Anderson MW. DNA adducts as a dosimeter for risk estimation. Environ. Health Perspect., 76: 3–8, 1978.

Bermudez E, Allen PF. The assessment of DNA damage and repair in rat nasal epithelial cells. Carcinogenesis, 5(11):1453–8, 1984.

Braga LMGM; Majerowicz J, Passos LAC, Gillioli R, Mattaraia VGM, Guaraldo AM, Diaz BL, Ko GM, Taricano, ID, Frajblay M, Stephano MA, Nascimento N, Massironi SMG, Lapchik VBV. Roedores e lagomorfos mantidos em instalações de instituições de Ensino ou pesquisa científica. In: Guia Brasileiro de produção manutenção ou utilização de animais em atividades de ensino ou pesquisa científica. Fascículo 2, 1ª Edição. Brasília: Ministério da Ciência, Tecnologia e Inovações e Comunicações, 2019. ISBN: 978-85-88063-76-1

Cai Y, Cao H, Wang F, Zhang Y, Kapranov P. Complex genomic patterns of abasic sites in mammalian DNA revealed by a high-resolution SSiNGLe-AP method. Nature Comm., 13: 5868 – 5888, 2022.

Camargo AP, Nakahara TS, Firmino LER, Netto PHM, do Nascimento JBP, Donnard ER, Galante PAF, Carazzolle MF, Malnic B, Papes F. Uncovering the mouse olfactory long non-coding transcriptome with a novel machine-learning model. DNA Res.; 26(4): 365–378, 2019.

Canugovi C, Misiak M, Scheibye-Knudsen M, Croteau DL, Mattson MP, Bohr VA. Loss of NEIL1 causes defects in olfactory function in mice. Neurobiology of Aging, v. 36, n. 2, p. 1007–1012, 2015.

Chen Y, Lun AT, Smyth GK. 2016. From reads to genes to pathways: Differential expression analysis of rna-seq experiments using rsubread and the edger quasi-likelihood pipeline. F1000Res. 5:1438.

Cheung MC, Jang W, Schwob JE, Wachowiak M. Functional recovery of odor representations in regenerated sensory inputs to the olfactory bulb. Frontiers in Neural Circuits. 7:207 1–16; 2014.

Cone CD Jr., Cone CM. Induction of mitosis in mature neurons in central nervous system by sustained depolarization. Nature. 192: 155–157; 1976.

Dan X, Wechter N, Gray S, Mohanty JG, Croteau DL, Bohr VA. Olfactory dysfunction in aging and neurodegenerative diseases. Ageing Res. Rev., 70: 101416, 2021.

Davis AJ, Chen DJ. DNA double strand break repair via non-homologous end-joining. Translational Cancer Research. 2(3): 130–143; 2013.

Doetsch F, Caillé I, Lim DA, García-Verdugo JM, Alvarez-Buylla A. Subventricular Zone Astrocytes Are Neural Stem Cells in the Adult Mammalian Brain. Cell, v. 97, p. 703–716, 1999.

Durinck S, Moreau Y, Kasprzyk A, Davis S, De Moor B, Brazma A, Huber W. 2005. Biomart and bioconductor: A powerful link between biological databases and microarray data analysis. Bioinformatics. 21(16):3439–3440.

Durinck S, Spellman PT, Birney E, Huber W. 2009. Mapping identifiers for the integration of genomic datasets with the r/bioconductor package biomart. Nat Protoc. 4(8):1184–1191.

Eriksson PS, Perfilieva E, Björk-Eriksson T, Alborn AM, Nordborg C, Peterson DA, Gage FH. Neurogenesis in the adult human hippocampus. Nature Med., 4(11): 1313–7, 1998.

Ge SX, Jung D, Yao R. 2020. Shinygo: A graphical gene-set enrichment tool for animals and plants. Bioinformatics. 36(8):2628–2629.

Genschel J, Modrich P. Mechanism of 5’-directed excision in human mismatch repair. Mol. Cell 12:1077–1086, 2003.

Hamilton M, Van Remmen H, Drake JA, Yang H, Guo ZM, Kewitt K, Walter CA, Richardson A. Does oxidative damage to DNA increase with age? Proc. Natl. Acad. Sci. USA, 98 (18) 10469–10474, 2001.

Hanchate NK, Kondoh K, Lu Z, Kuang D, Ye X, Qiu X, Pachter L, Trapnell C, Buck LB. Single-cell transcriptomics reveals receptor transformations during olfactory neurogenesis. Science, 350(6265): 1251–5, 2015.

Ishii T, Omura M, Mombaerts P. Protocols for two- and three-color fluorescent RNA in situ hybridization of the main and accessory olfactory epithelia in mouse. Journal of neurocytology. 33(6): 657–69; 2004.

Kebir S, Hattingen E, Niessen M, Rauschenbach L, Fimmers R, Hummel T, Schäfer N, Lazaridis L, Kleinschnitz C, Herrlinger U, Scheffler B, Glas M. Olfactory function as an independent prognostic factor in glioblastoma. Neurology, 94(5): e529–e537, 2020.

Kempermann G, Gage FH, Aigner L, et al. Human adult neurogenesis: evidence and remaining questions. Cell Stem Cell, 23(1):25–30, 2018.

Kim D, Pertea G, Trapnell C, Pimentel H, Kelley R, Salzberg SL. 2013. Tophat2: Accurate alignment of transcriptomes in the presence of insertions, deletions and gene fusions. Genome Biol. 14(4): R36.

Kimura KC, Fukumasu H, Chaible LM, Lima CE, Horst MA, Matsuzaki P, Sanches DS, Pires CG, Silva TC, Pereira TC, Mello ML, Matera JM, Dias RA, Monnereau A, Sasco AJ, Saldiva PH, Dagli ML. Evaluation of DNA damage by the alkaline comet assay of the olfactory and respiratory epithelia of dogs from the city of São Paulo, Brazil. Exp. Toxicol. Pathol., 62(3): 209–119, 2010.

Klein JA, Longo-Guess CM, Rossmann MP, Seburn KL, Hurd RE, Frankel WN, Bronson RT, Ackerman SL. The harlequin mouse mutation downregulates apoptosis-inducing factor. Nature, 419 (6905): 367–74, 2002.

Langmead B, Salzberg SL. 2012. Fast gapped-read alignment with bowtie 2. Nat Methods. 9(4):357–359.

Liao Y, Smyth GK, Shi W. The subread aligner: Fast, accurate and scalable read mapping by seed-and-vote. Nucleic Acids Res. 41(10):e108, 2013.

Liao Y, Smyth GK, Shi W. Featurecounts: An efficient general purpose program for assigning sequence reads to genomic features. Bioinformatics. 30(7):923–930, 2014.

Lois C, Alvarez-Buyiia A. Long-Distance Neuronal Migration in the Adult Mammalian Brain. Science, 264(5162), 1145–1148. 1994.

Mani C, Reddy OH, Palle K. DNA repair fidelity in stem cell maintenance, health, and disease. Biochimica et Biophysica Act - Molecular Basis of Disease, 1866(4): 165444, 2020.

Marteijn JA, Lans H, Vermeulen W, Hoeijmakers JH. Understanding nucleotide excision repair and its roles in cancer and ageing. Nature Reviews Molecular Cell Biology, v. 15, n. 7, p. 465–481, 2014.

Maynard S, Swistowska AM, Lee JW, Liu Y, Liu ST, Da Cruz AB, Rao M, de Souza-Pinto NC, Zeng X, Bohr VA. Human embryonic stem cells have enhanced repair of multiple forms of DNA damage. Stem Cells, 26(9):2266–74, 2008.

McCarthy DJ, Chen Y, Smyth GK. Differential expression analysis of multifactor rna-seq experiments with respect to biological variation. Nucleic Acids Res. 40(10):4288–4297, 2012.

Misiak M, Vergara Greeno R, Baptiste BA, Sykora P, Liu D, Cordonnier S, Fang EF, Croteau DL, Mattson MP, Bohr VA. DNA polymerase β decrement triggers death of olfactory bulb cells and impairs olfaction in a mouse model of Alzheimer’s disease. Aging Cell, v. 16, n. 1, p. 162–172, 2017.

Modrich, P. Mechanisms in E. coli and Human Mismatch Repair (Nobel Lecture). Review Angew Chem Int Ed Engl., 55(30):8490–501, 2016.

Parma V, Ohla K, GCCR Group, et al. More than smell-COVID-19 is associated with severe impairment of smell, taste, and chemesthesis. Chem Senses, 45(7):609–622, 2020.

Perry G, Castellani RJ, Smith MA, Harris PL, Kubat Z, Ghanbari K, Jones PK, Cordone G, Tabaton M, Wolozin B, Ghanbari H. Oxidative damage in the olfactory system in Alzheimer’s disease. Acta Neuropathologica, 106(6): 552– 556, 2003.

Pffafl, MW. A new mathematical model for relative quantification in real-time RT-PCR. Nucleic Acids Research. 1;29(9): e45, 2001.

Qiu X, Hill A, Packer J, Lin D, Ma Y-A, Trapnell C. Single-cell mRNA quantification and differential analysis with census. Nature Methods. 14(3):309–315, 2017

Qiu X, Mao Q, Tang Y, Wang L, Chawla R, Pliner HA, Trapnell C. Reversed graph embedding resolves complex single-cell trajectories. Nature Methods. 14(10):979–982, 2017.

Ren J, Wang X, Dong C, Wang G, Zhang W, Cai C, Qian M, Yang D, Ling B, Ning K, Mao Z, Liu B, Wang T, Xiong L, Wang W, Liang A, Gao Z, Xu J. Sirt1 Protects Subventricular Zone-Derived Neural Stem Cells from DNA Double-Strand Breaks and Contributes to Olfactory Function Maintenance in Aging Mice. Stem Cells, 40(5):493–507, 2022.

Eriksson PS, Perfilieva E, Björk-Eriksson T, Alborn AM, Nordborg C, Peterson DA, Gage FH. Neurogenesis in the adult human hippocampus. v. 4, n. 11, p. 1313–1317, 1998.

Robinson MD, McCarthy DJ, Smyth GK. Edger: A bioconductor package for differential expression analysis of digital gene expression data. Bioinformatics. 26(1):139–140, 2010.

Schwob, J. Neural Regeneration and the Peripheral Olfactory System. The Anatomical Record, 269:33–49; 2002.

Sherman MH, Bassing CH, Teitell MA. DNA damage response regulates cell differentiation Trends Cell Biol., 21(5): 312–319, 2011.

Sjakste N, Riekstiņa U. DNA damage and repair in the differentiation of stem cells and cells of connective cell lineages: A trigger or a complication? Eur J Histochem., 65(2): 3236, 2021.

Souza-Pinto NC, Wilson III DM, Stevnsner TV, Bohr VA. Mitochondrial DNA, base excision repair and neurodegeneration. DNA repair, 7: 1098–1109. 2008.

Sykora P, Yang J-L, Ferrarelli LK, Tian J, Tadokoro T, Kulkarni A, Weissman L, Keijzers G, WilsonIII DM, Mattson MP, Bohr VA. Modulation of DNA base excision repair during neuronal differentiation Neurobiol. Aging, 34(7): 1717– 1727, 2012.

Tang H, Chen L, Dai Z, Zhang W, Wang T, Wu L, Wang G, Bian P. Enhancement of DNA damage repair potential in germ cells of Caenorhabditis elegans by a volatile signal from their irradiated partners. DNA Repair, 86: 2, 2020.

Team RC. 2022. R: A language and environment for statistical computing.

Tofilon PJ, Meyn RE. Influence of Cellular Differentiation on Repair of Ultraviolet-Induced DNA Damage in Murine Proadipocytes. Radiation Research, 116(2): 217–227, 1988.

Trapnell C, Cacchiarelli D, Grimsby J, Pokharel P, Li S, Morse M, Lennon NJ, Livak KJ, Mikkelsen TS, Rinn JL. The dynamics and regulators of cell fate decisions are revealed by pseudotemporal ordering of single cells. Nature Biotechnology. 32(4):381–386, 2014.

Vaishnav RA, Getchell ML, Huang L, Hersh MA, Stromberg AJ, Getchell TV. Cellular and molecular characterization of oxidative stress in olfactory epithelium of Harlequin mutant mouse. J. Neurosci. Res., 86(1): 165–182, 2008.

Valavanidis A, Vlachogianni T, Fiotakis K, Loridas S. Pulmonary oxidative stress, inflammation and cancer: Respirable particulate matter, fibrous dusts and ozone as major causes of lung carcinogenesis through reactive oxygen species mechanisms. Int. J. of Environ. Res. and Public Health, 10(9): 3886– 3907, 2013.

Wang X, Spandidos A, Wang H, Seed B. PrimerBank: a PCR primer database for quantitative gene expression analysis, 2012 update. Nucleic Acids Res., 40: D1144–9, 2012.

Zhang S, Wang M, Villalta PW, Lindgren BR, Upadhyaya P, Lao Y, Hecht SS. Analysis of pyridyloxobutyl and pyridylhydroxybutyl DNA adducts in extrahepatic tissues of F344 rats treated chronically with 4-(methylnitrosamino)-1-(3-pyridyl)-1-butanone and enantiomers of 4-(methylnitrosamino)-1-(3-pyridyl)-1-butanol. Chem. Res. Toxicol., 22(5): 926–936, 2009.

