## Supplemental tables for "Expression of DNA repair genes is modulated during differentiation of olfactory sensory neurons"

**Supplemental Material**

**Table S1**: Primer sequences for long extension PCR

| Amplicon | Length (bp) | Primer sequence |
| --- | --- | --- |
| Mitochondrial long *forward* | 10.000 | GCCAGCCTGACCCATAGCCATAATAT |
| Mitochondrial long *reverse* |  | GAGAGATTTTATGGGTGTAATGCGG |
| Mitochondrial short (RNR1)  *forward* | 128 | AACCTCCATAGACCGGTGTAAAA |
| Mitochondrial short (RNR1) *reverse* |  | TTTATCACTGCTGAGTCCCGT |
| Nuclear long (β-globin)  *forward* | 8.700 | TTGAGACTGTGATTGGCAATGCCT |
| Nuclear long (β-globin)  *reverse* |  | GCCTGGACTTTGCCCCTAAT |
| Nuclear short (HPRT)  *forward* | 134 | GCCTGGACTTTGCCCCTAAT |
| Nuclear short (HPRT)  *reverse* |  | CGCCTTTCCACTCTTCAGGT |

**Table S2**: Experimental conditions for XL-PRC

| Amplicon | DNA (ng) | Cycling conditions |
| --- | --- | --- |
| Nuclear short | 120 | 94°C – 3min (1 cycle)  94°C – 45s; 56°C – 30s; 72°C – 1min (22 cycles) |
| Nuclear long | 30 | 94°C – 30s (1 cycle)  94°C – 30s; 62°C – 30s; 68°C – 15min (33 cycles) |
| Mitochondrial short | 3 | 94°C – 3min (1 cycle)  94°C – 45s; 56°C – 30s; 72°C – 1min (22 cycles) |
| Mitochondrial long | 6 | 94°C – 30s (1 cycle)  94°C – 30s; 56°C – 30s; 68°C – 18min (28 cycles) |

**Table S3:** Primer sequences were obtained by *PimerBank* (Wang, 2012), and validated by BLAST (<https://blast.ncbi.nlm.nih.gov/Blast.cgi>).

| Pathway | Gene | Forward | Reverse |
| --- | --- | --- | --- |
| BER | UNG | TTCGGGAAGCCGTACTTCG | CATCTGGGTCCATGTGAACAC |
|  | POLB | TGAACCATCATCAACGAATTGGG | CCATGTCTCCACTCGACTCTG |
|  | OGG1 | CTGCCTAGCAGCATGAGACAT | CAGTGTCCATACTTGATCTGCC |
|  | APEX1 | ACGGGGAAGAACCCAAGTC | TCCACATTCCAGGAGCATATCT |
| HR | Rad 52 | CTTTGTTGGTGGGAAGTCTGT | CGGCTGCTAATGTACTCTGGAC |
|  | Exo1 | TGGCTGTGGATACCTACTGTT | ATCGGCTTGACCCCATAAGAC |
|  | BRCA1 | CGAATCTGAGTCCCCTAAAGAGC | AAGCAACTTGACCTTGGGGTA |
|  | BRCA2 | ATGCCCGTTGAATACAAAAGGA | ACCGTGGGGCTTATACTCAGA |
| NER | XPC | TCCAGGGGACCCCACAAAT | GCTTTTTGGGTGTTTCTTTGCC |
|  | XPA | CCCAAAATGATTGACACCAAAGG | TGGTTCATAAGGTACGAGTCCA |
|  | ERCC6 | GAGACCACAGAGAAACGTCCA | CTCCTGCTCACTGGTTTGCT |
|  | POLD1 | TCGCAGTTTGAGGCGAACC | CTGAGCCCACATAGTGGTCA |
| NHEJ | XRCC6 | ATGTCAGAGTGGGAGTCCTAC | TCGCTGCTTATGATCTTACTGGT |
|  | XRCC5 | ATGGCGTGGTCCGGTAATAAG | CCTGTCGTTGGACAAACATAGTC |
|  | PRKDC | AAACCTGTTCCGAGCTTTTCTG | TAAGGGCACAGCATATCGCTT |
|  | LIG4 | TTTGCAGACTTATGTTCCACACT | TCCTTTCTGTTCTTATGAAGGGC |
| Loading control | HMBS | AAGGGCTTTTCTGAGGCACC | AGTTGCCCATCTTTCATCACTG |
| Olfactory markers | Ric-8B | AGCTGGTTCGTCTCATGACAC | CAGCGTTCCCATAGCCAGTG |
|  | Ngn1 | CCAGCGACACTGAGTCCTG | CGGGCCATAGGTGAAGTCTT |
|  | OMP | TCCGTCTACCGCCTCGATTT | CGTCTGCCTCATTCCAATCCA |

**Table S4**: Efficiency values were obtained by *qPCR Efficiency Calculator* (Thermo Fisher Scientific), available at <https://www.thermofisher.com/br/en/home/brands/thermo-scientific/molecular-biology/molecular-biology-learning-center/molecular-biology-resource-library/thermo-scientific-web-tools/qpcr-efficiency-calculator.html>

|  | Gene | Amplification factor |  | Gene | Amplification factor |
| --- | --- | --- | --- | --- | --- |
| BER | UNG | 2,15 | **NER** | XPC | 2,09 |
|  | POLB | 2,04 |  | XPA | 2,09 |
|  | OGG1 | 1,96 |  | ERCC6 | 2,06 |
|  | APEX1 | 1,97 |  | POLD1 | 2,05 |
| HR | BRCA1 | 1,97 | **NHEJ** | XRCC6 | 1,99 |
|  | BRCA2 | 1,97 |  | XRCC5 | 1,91 |
|  | RAD52 | 2,00 |  | PRKDC | 2,13 |
|  | EXO1 | 1,99 |  | LIG4 | 2,12 |

|  | Gene | Amplification factor |
| --- | --- | --- |
| Olfactory markers | OMP | 2,16 |
|  | NGN1 | 1,98 |
|  | RIC8B | 1,97 |
| Normalizer | HMBS | 2,11 |

**Table S5**: Primer sequence and insert size for the RNA probe used in in situ hybridization.

| Target | Amplicon (bp) | Sequence |
| --- | --- | --- |
| APE1 *forward* | 995 | ACGGGGAAGAACCCAAGTC |
| APE1 *reverse* |  | AACACCTGAAGGCTAAAACACCAG |

**Table S6**: Expression trajectory analysis of 481 DNA repair genes from single-cell gene expression data of Hanchate et al (2015) with Monocle.

The complete list is available in the Excel file included with the supplemental material.
